## supporting information for "Separate-scan atomic force microscope for fast infrared scattering-type scanning near-field optical microscope"

To calculate critical angle, refractive index at wavelength 6 μm for Si 3.47^1^, water 1.26^2^, ZnSe 2.43^3^ are used.

Parameter for FDTD calculations by Lumerical. 200 fs simulation time, 300 K simulation temperature, source shape Gaussian, injection axis z-axis, 5000 nm waist radius, center frequency 1600 cm^-1^, frequency width 500 cm^-1^.

The Si prism, the sSNOM probe are mimicked by the Si half sphere, 35 nm radius Platinum (Pt) sphere and Pt pyramid, respectively.

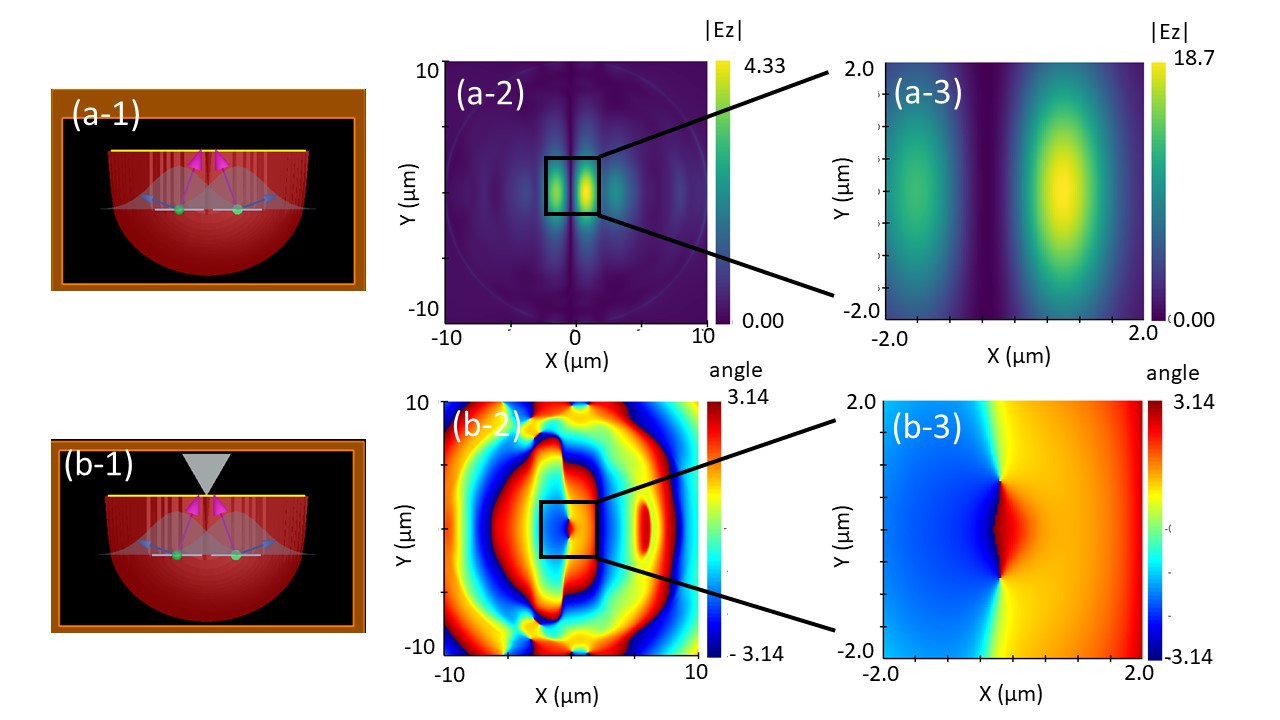

Supp. Fig. 1 FDTD to show how E-field propagate on Si-air interface from two gaussian sources. (a-1) Side view. 20° θ angle to have same focal point on the surface. (a-2) 6 µm of wavelength. Depending on wavelength, FDTD shows the asymmetrical amplitude.

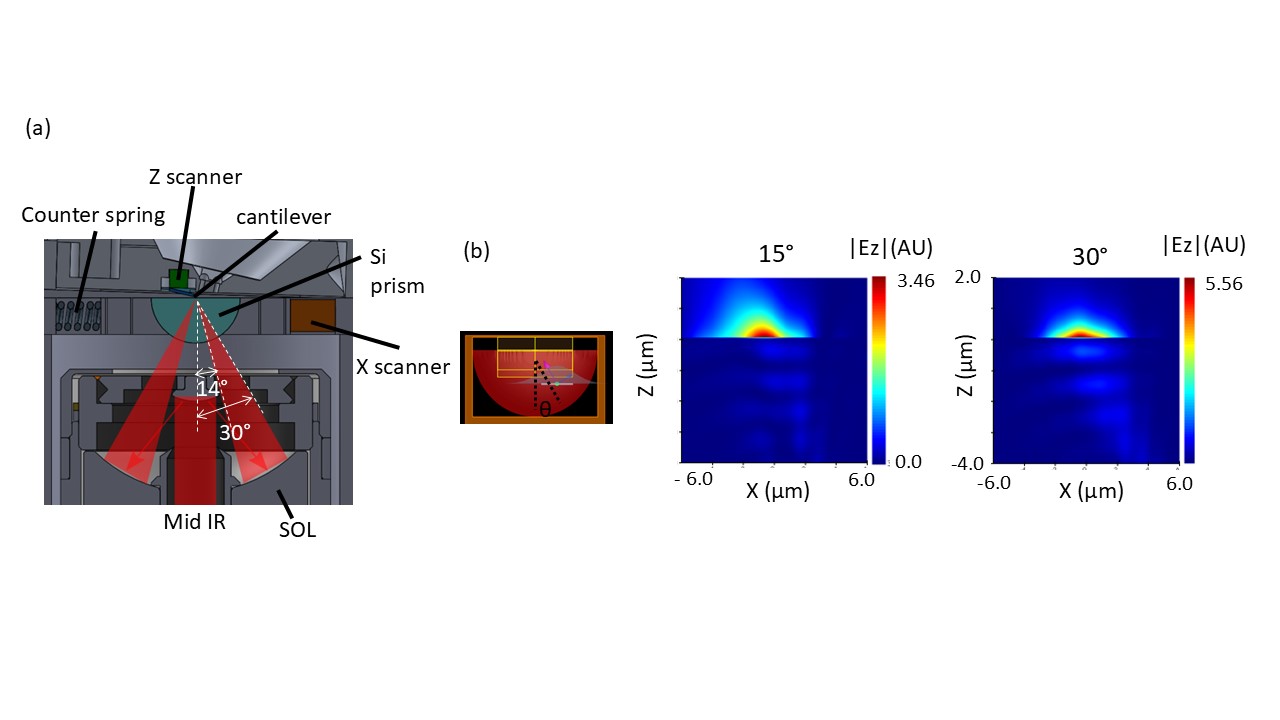

Supp. Fig. 2 Comparison of p-polarization of the electric field with different incident light angles. (a) Section view of BI-sSNOM. (b) FDTD calculation of the E-filed. Incident light angle 15° and 30°. Top row; E-field propagation around interface of Si and air. 30 nm mesh size.

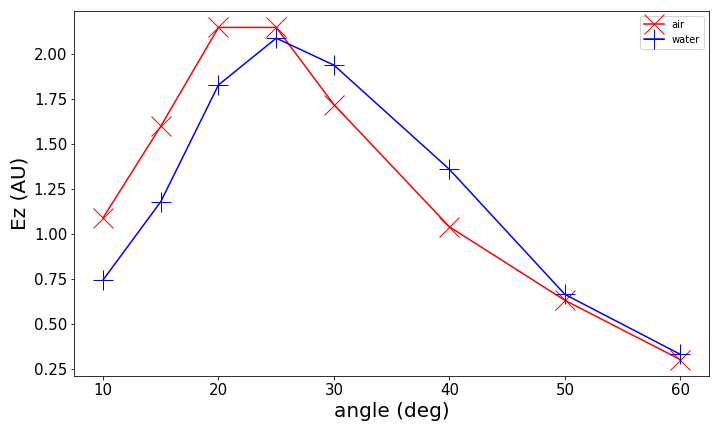

Supp. Fig. 3 FDTD calculations of E-field on the interface with different incident angle. Red, Maximum of Z component of E-field in air. Blue, Z component of E-field in water.

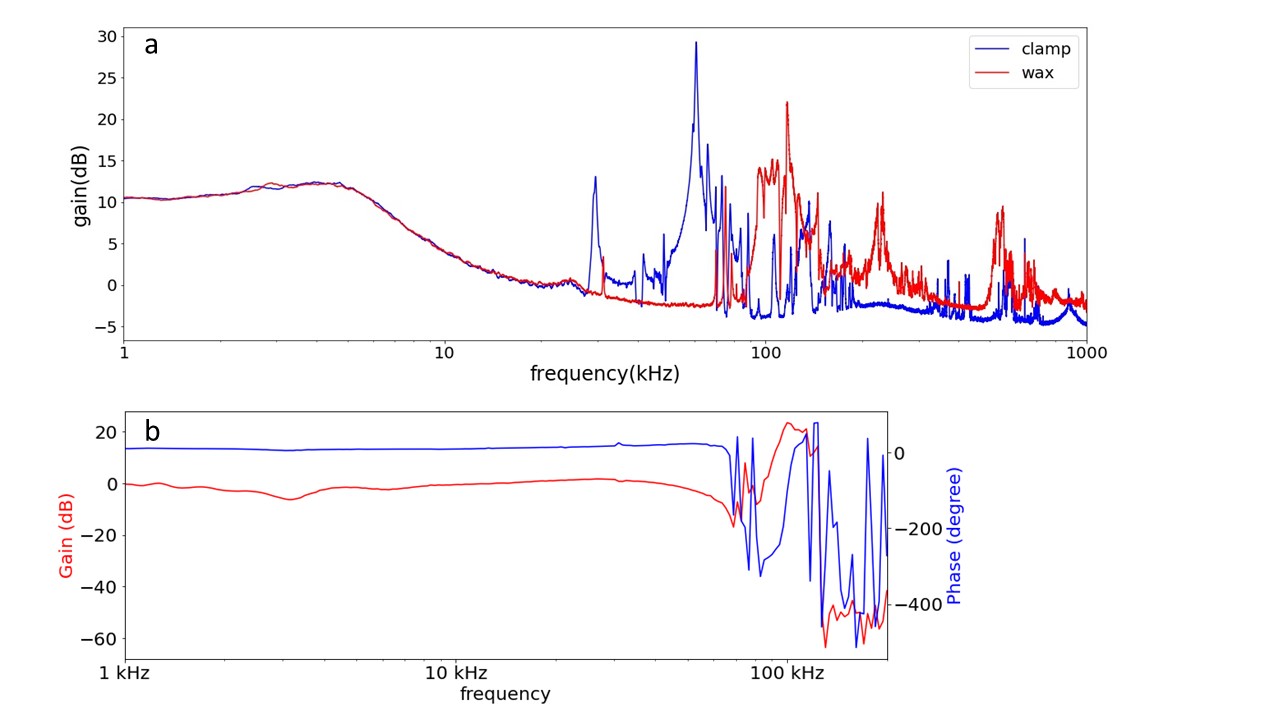

Supp. Fig. 4 Gain and phase plot of frequency response of Z-scanner. (a) Amplitude sweep using a SmarAct vibrometer, gain is calculated by a python code after the measurement. (b) Amplitude and phase sweep measured using a custom-built interferometer. Cantilever is mounted with wax. The laser spot size is larger than cantilever body and the ambient vibration isolation is imperfect for the custom-built interferometer.

Gain is calculated following, Gain (dB) = 20log_10_(V_out_/V_baseline_). Line plot of wax in (a) and gain plot in (b) are essentially same.

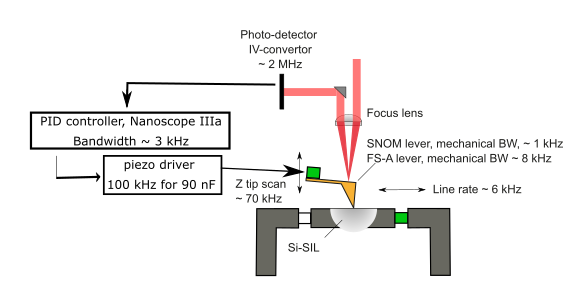

Supp. Fig. 5 Schematic of Feedback loop for the AFM in air. Bandwidth of main components.

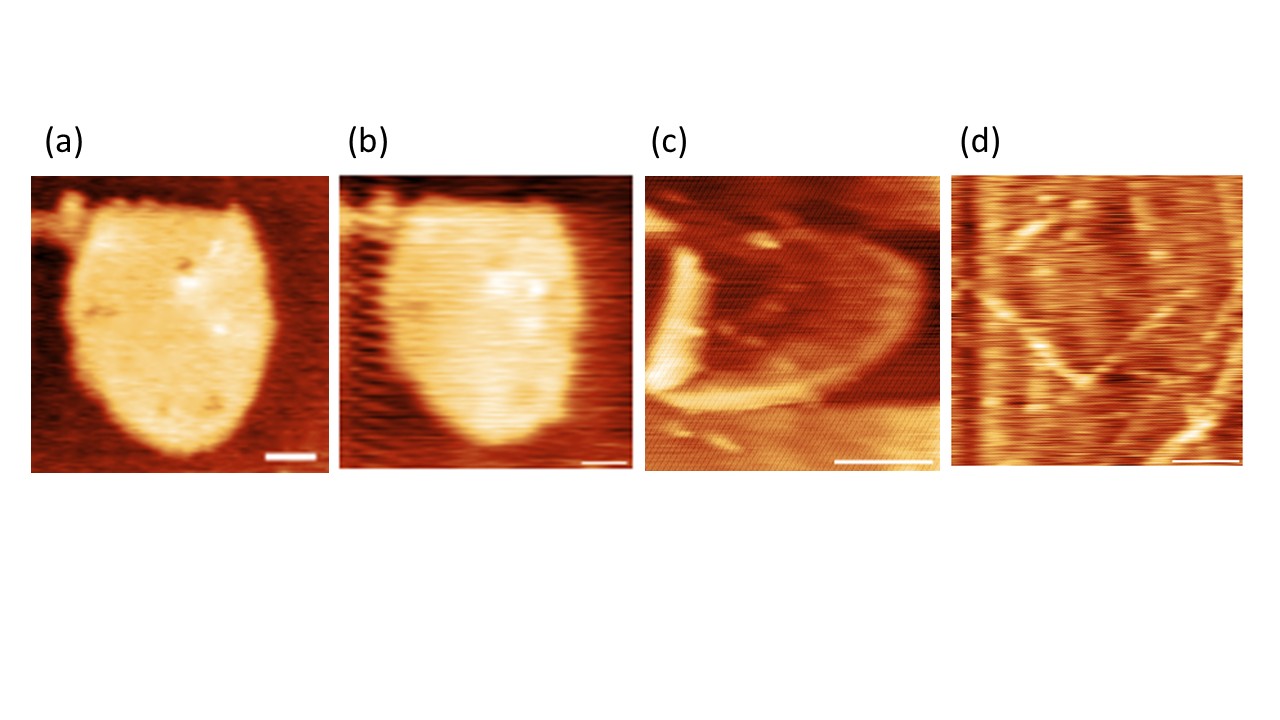

Supp. Fig. 6 AFM image with different cantilevers. All scale bar 200 nm. (a,b) PM in buffer by fast-scan C lever. (a)60 lines/s. (b)118 lines/s. (c) PM in air by fast-scan A with photothermal tapping. 256 pixels/line, 118 lines/s. (d) Actin filament in air by ultra-short lever. 118 lines/s. Data are treated by tilt compensation and 2 degree of Polynomial filter by Gwyddion

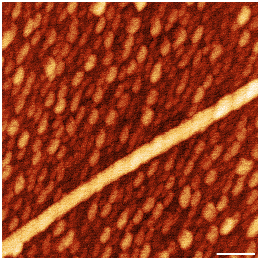

Supp. Fig. 7, AFM image of microtubule. Photothermal tapping in air with fast-scan A lever. Scale bar 200 nm. 2 lines/s.

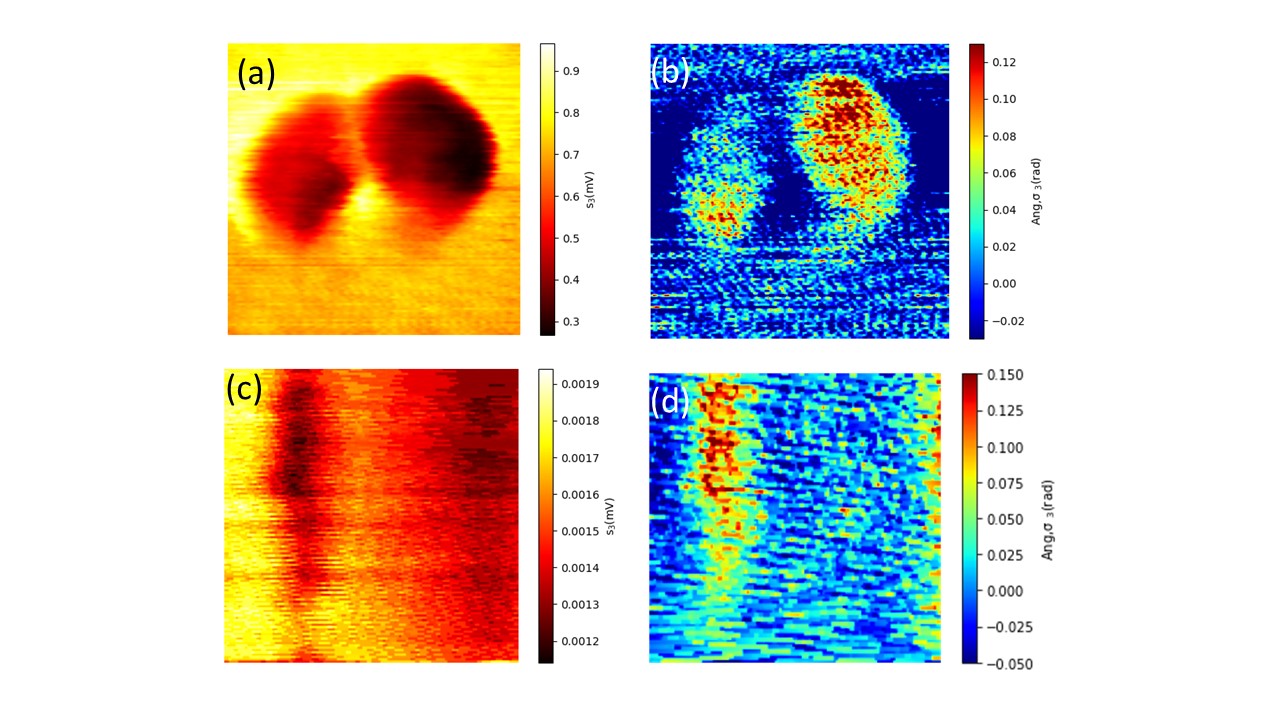

Supp. Fig. 8, Optical images without median filter which are shown in Fig. 9 and Fig. 10.

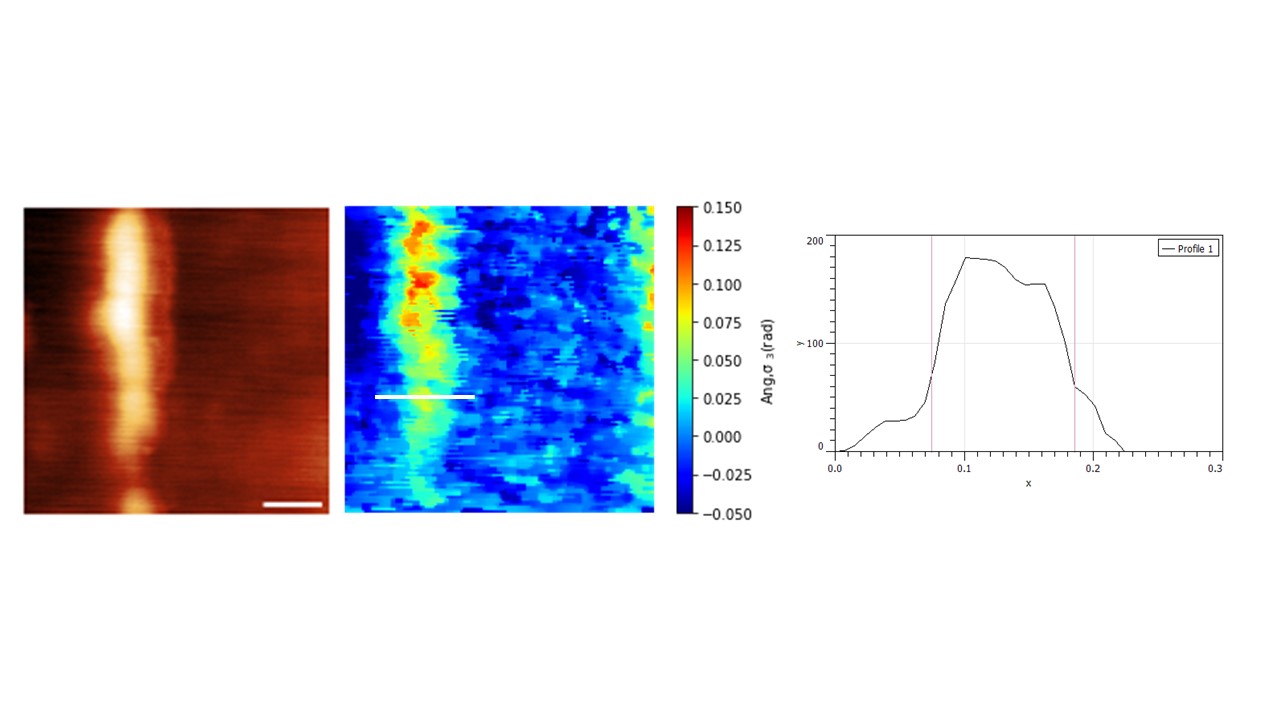

Supp. Fig. 9, The aspect ratio of optical phase image. Cross-section shows 110 nm. The aspect ratio of sSNOM probe is calculated as follow. (110 nm – 25 nm (outer diameter of microtubule))/2 = 42.5 nm.

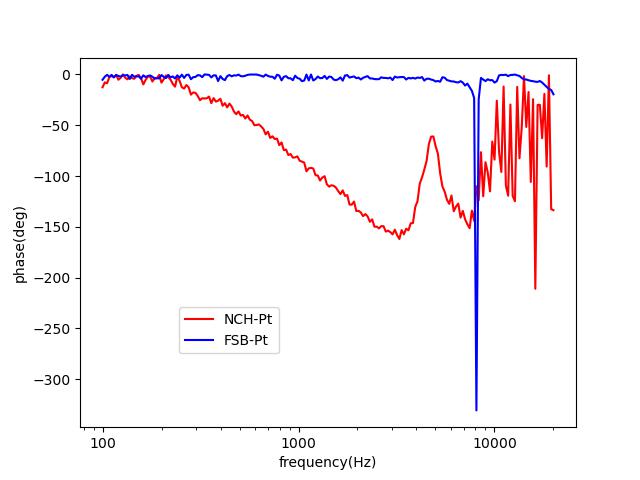

Supp. Fig. 10, Mechanical bandwidth of sSNOM levers in water. While the tip and the sample are engaged with the tapping mode at the resonance frequency, the response of the cantilever to actuation of the Z-scanner were measured at each frequency. The phase decay starts at ca 250 Hz and ca 8 kHz for Arrow-NCHPt and FSB-Pt, respectively. Oscillation amplitudes of the cantilever for optimal near-field signal or high-speed AFM are above 50 nm or ca 2 nm, respectively. This difference gives a discrepancy between the measurement value and the theoretical value of the mechanical bandwidth. FSB-Pt has a mechanical bandwidth more than an order of magnitude higher than conventional sSNOM levers in water.

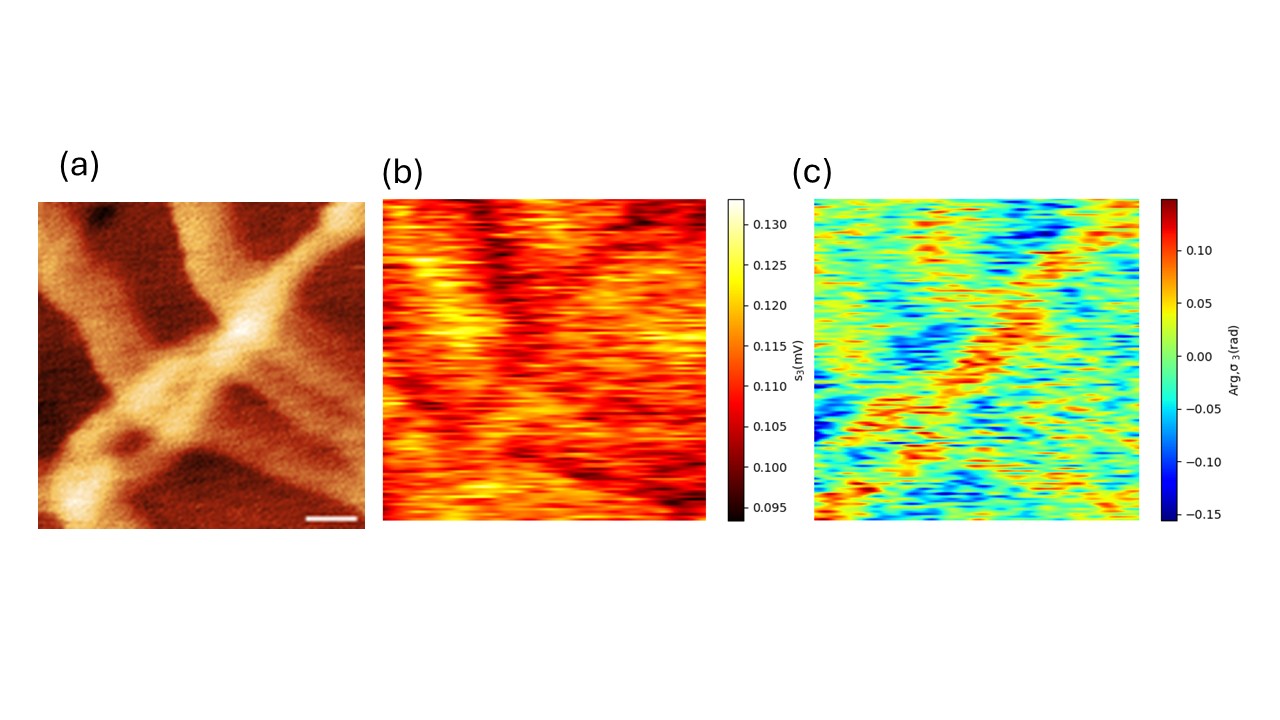

Supp. Fig. 11, sSNOM experiment of microtubules in tubulin buffer. Without median filter. Wavenumber 1665 cm^-1^. 9.86 lines/s (a) AFM topography. Scale bar 200 nm. (b) Optical amplitude image. (c) Optical phase image.

1.	Shkondin E, Takayama O, Panah MEA, et al. Large-scale high aspect ratio Al-doped ZnO nanopillars arrays as anisotropic metamaterials. *Opt Mater Express*. 2017;7(5):1606-1627. doi:10.1364/OME.7.001606

2.	Hale GM, Querry MR. Optical Constants of Water in the 200-nm to 200-$μ$m Wavelength Region. *Appl Opt*. 1973;12(3):555-563. doi:10.1364/AO.12.000555

3.	Connolly J, diBenedetto B, Donadio R. Specifications Of Raytran Material. In: *Proc.SPIE*. Vol 0181. ; 1979:141-144. doi:10.1117/12.957359
